# *Wisteria floribunda* agglutinin enhances *Zaire ebolavirus* entry through interactions at specific N-linked glycosylation sites on the virus glycoprotein complex

**DOI:** 10.1101/2024.12.05.626977

**Authors:** Joshua D. Duncan, Monika Pathak, Barnabas J. King, Holly Bamber, Paul Radford, Jayasree Dey, Charlotte Richardson, Stuart Astbury, C. Patrick McClure, Jonathan K. Ball, Richard A. Urbanowicz, Alexander W. Tarr

## Abstract

Entry of *Zaire* e*bolavirus* (EBOV) into a host cell is a complex process requiring interactions between the viral glycoproteins (GP) and cellular factors. These entry factors are cell-specific and can include cell surface lectins and phosphatidylserine receptors. NPC1 is critical to the late stage of the entry process. Entry has been demonstrated to be enhanced by interactions between the virion and surface-expressed lectins, which interact with carbohydrate moieties attached to the GP. In addition, soluble lectins, including mannose binding lectin (MBL), can enhance entry *in vitro*. However, the mechanism of lectin-mediated enhancement remains to be defined.

This study investigated the potential of three plant lectins, *Wisteria floribunda* agglutinin (WFA), soybean agglutinin (SBA) and *Galanthus nivalis* agglutinin (GNA), which possess different carbohydrate binding specificities, to enhance EBOV entry by binding to the GP. WFA was observed to potently enhance entry of lentiviral pseudotype viruses (PVs) expressing the GP of three *Ebolavirus* species (Zaire, Sudan [SUDV] and Reston [RESTV]), with the greatest impact on EBOV. SBA had a modest enhancing effect on entry that was specific to EBOV, while GNA had no impact on entry of any of the *Ebolavirus* species. None of the lectins enhanced entry of control PVs expressing the surface proteins of other RNA viruses tested. WFA was demonstrated to bind directly with the EBOV-GP via the glycans, and mutational analysis implicated N^238^ as contributing to the interaction. Furthermore, enhancement was observed in both human and bat cell lines indicating a highly conserved mechanism of action. We conclude that binding of WFA to EBOV GP through interactions including the glycan at N^238^ results in GP alterations that enhance entry, providing evidence of a mechanism for lectin-mediated virus entry enhancement. Targeting lectin-ligand interactions presents a potential strategy for restricting *Ebolavirus* entry.

## Introduction

*Zaire ebolavirus* (EBOV) is a negative-strand RNA virus that represents a significant threat to human health. EBOV infection causes haemorrhagic fever^1^, resulting in a case fatality rate of 60-90%^2^ . Since its discovery in 1976, EBOV has been associated with multiple outbreaks in central Africa, the largest of which being the 2018-20 North Kivu epidemic in the Democratic Republic of the Congo, and the 2013-16 epidemic in West Africa which resulted in 2,200 and 11,000 deaths, respectively^3,4^. These outbreaks are believed to result from spillover events originating from an as-yet unidentified animal reservoir(s).

EBOV encodes an envelope glycoprotein (GP) complex that is incorporated onto the surface of the filamentous virion. GP comprises covalently linked heterodimers of GP_1_ and GP_2_ subunits and acts as a Class I fusion protein, mediating host cell entry via cellular attachment and membrane fusion within late-stage endosomes^5–7^. The N-terminus possesses the GP_1_ subunit, which possesses a glycan cap that masks the GP_2_ subunit that contains the fusion peptide^7,8^. EBOV entry requires cleavage of the GP in the endosome to remove the glycan cap^9^, exposing the binding site for the cholesterol transporter protein, Niemann-Pick type C1 (NPC1)^10,11^.This binding site is located in a pocket in the GP_1_ molecule, involving amino acids T^83^, I^113^ and L^122 11^.

During maturation the EBOV-GP undergoes extensive modification via *N*-, *O*- and *C*-linked glycosylation, leading to the occlusion of epitopes from recognition by host antibodies^12^ and receptor attachment^13^. Removal of the carbohydrate from the GP complex is an essential step in entry. A total of 17 predicted *N*-linked glycosylation sites exist in Zaire EBOV-GP, 15 of which are in the GP_1_ subunit and predominantly located within the glycan cap and mucin-like domain (MLD) regions. Glycan profiling has shown that when expressed in human cells, these sites are occupied by simple mannose or complex oligosaccharides, indicating that processing of the EBOV-GP occurs in the Golgi apparatus^13^. The glycosylation of GP is complex, with varying sugars present at specific *N*-linked sites, and substantial *O*-linked glycosylation occurring in the mucin-like domain (MLD)^13^. One site in the glycan cap (N^257^) has been found to have a restricted, mannose-type sugar, in contrast to the complex glycosylation exhibited at other sites^13^.

The extensive glycosylation of the GP facilitates EBOV entry^14^. Interaction with lectins has been proposed as potential attachment receptors^15^, but evidence of their role as definitive entry receptors is lacking^16^. Lectins are a diverse group of proteins that possess a range of different carbohydrate binding specificities. In humans they contribute to innate immunity by sensing pathogen-associated molecular patterns (PAMPs)^17,18^. Interactions between lectins and viruses can have differing consequences. Soluble lectins can effectively neutralise virus particles through direct interactions with virus glycoproteins^19–24^. Conversely, transmembrane lectins, such as human macrophage C-type lectin and dendritic cell- or liver/ lymph node-specific ICAM3-grabbing non-integrin (DC-SIGN; CD209), CLEC4M/L-SIGN (CD299), human macrophage lectin specific for galactose/N-acetylgalactosamine (hMGL), liver and lymph node sinusoidal endothelial cell C-type lectin (LSECtin) and asialoglycoprotein receptor I (ASGPRI) all enhance *in vitro* EBOV entry in specific cell-types^25,26^. However, the C-type lectins are not essential for entry, as evidenced by their negligible expression on some permissive cell types^27^, and their inability to confer susceptibility to non-permissive cells^16,28^.

In addition to membrane-associated lectins, secreted lectins such as mannose-binding lectin (MBL) have been demonstrated to enhance infection. MBL is a soluble component of the humoral innate immune response. Through recognition of pathogen-associated carbohydrate patterns, MBL can opsonise a target leading to neutralisation. MBL has previously been shown to neutralise a variety of viruses^18^. In the context of EBOV, MBL exhibits complement-dependent neutralising activity towards retroviral pseudotyped particles bearing authentic EBOV-GP^29^, and can protect against lethal challenge^30^. Strikingly, however, in the absence of complement, MBL enhances EBOV cellular entry^31^. This enhancement has been proposed to be a function of MBL directly coupling virus particles to cell surfaces, acting as a ‘bridging’ receptor and promoting uptake into endosomes^31,32^.

Plant-derived lectins, in addition to those expressed by animal cells, have been studied for their antiviral activity and their use in characterising the glycan profiles of viral glycoproteins through large lectin-based arrays^33^. Antiviral activity of plant lectins towards a wide range of viruses has been reported^34^. The mechanisms of antiviral activity by lectins vary, with some blocking viral replication and others neutralising virus entry into a host cell^35^. For example, BanLec, a lectin derived from *Musa acuminata*^36^, has previously been shown to inhibit both EBOV replication and entry^37^. As such, plant lectins have potential as inhibitors for virus infections. A broad-spectrum antiviral protein may have application in the public health response to emerging virus infections.

In this study we addressed the possibility that interactions between soluble lectins with EBOV-GP enhance infection without the requirement for a proposed cell-expressed lectin binding receptor. The ability of plant-derived lectins with different carbohydrate-binding specificities to enhance EBOV-GP-mediated transduction of permissive cells was assessed. Using a retroviral pseudotype entry model incorporating full-length authentic EBOV-GP proteins^38^, three plant-derived lectins (*Wisteria floribunda* agglutinin, WFA; Soybean agglutinin, SBA; and *Galanthus nivalis* agglutinin, GNA), characterised by different saccharide specificities, were compared. WFA has specificity for LacdiNAc (β-d-GalNAc-[1→4]-d-GlcNAc) sugars, with lesser affinity for GalNac^39^. SBA binds to multiantennary carbohydrates with terminal GalNAc or Gal residues^40^. In contrast, GNA binds to sugars possessing high-mannose type simple sugars^41^. The presence of WFA significantly enhanced EBOV PV entry. This effect was observed in EBOV-GP variants and related GP proteins from other *Filovirus* species. WFA binding also reduced the neutralizing activity of an anti-GP monoclonal antibody (KZ52).

## Results

### Disaccharide-specific lectins specifically enhance *Ebolavirus* infection

To investigate the consequences of the presence of plant derived lectins during viral entry, a retroviral pseudotype entry model was employed, incorporating full-length authentic viral GPs (Figure 1). Two reference strains of Zaire EBOV were initially assessed, the Makona C15 (EBOV-C15) and wild-type Mayinga 1976 (EBOV-May). These strains represent the initial variant that appeared in the 2014 epidemic, and the original isolate of EBOV. GPs representing two other *ebolavirus* species, *Sudan ebolavirus* (SUDV) and *Reston ebolavirus* (RESTV), were also assessed. The GPs of Zaire EBOV, SUDV and RESTV share conserved *N-*linked glycan sites at N^40^, N^204^, N^238^, N^259^, N^270^, N^298^, N^563^and N^618^ (Supplementary figure 1). In addition, pseudotypes possessing viral GPs recovered from other viruses with Class I fusion proteins (*lymphocytic choriomeningitis mammarenavirus* (LCMV)), Class III fusion proteins (*Indiana vesiculovirus*, G protein (VSV) and *Rabies lyssavirus* (RABV)), and novel class fusion proteins (hepatitis C virus (HCV) E1/E2)^42^, were included in this analysis. Each of these viral glycoproteins exhibit *N*-linked glycosylation (SUDV-GP possesses 12 sites, RESTV-GP possesses 14, LCMV-GP possesses 11, both RABV-G and VSV-G possess two and HCV E1/E2 GPs together possess 16).

**Figure 1.**
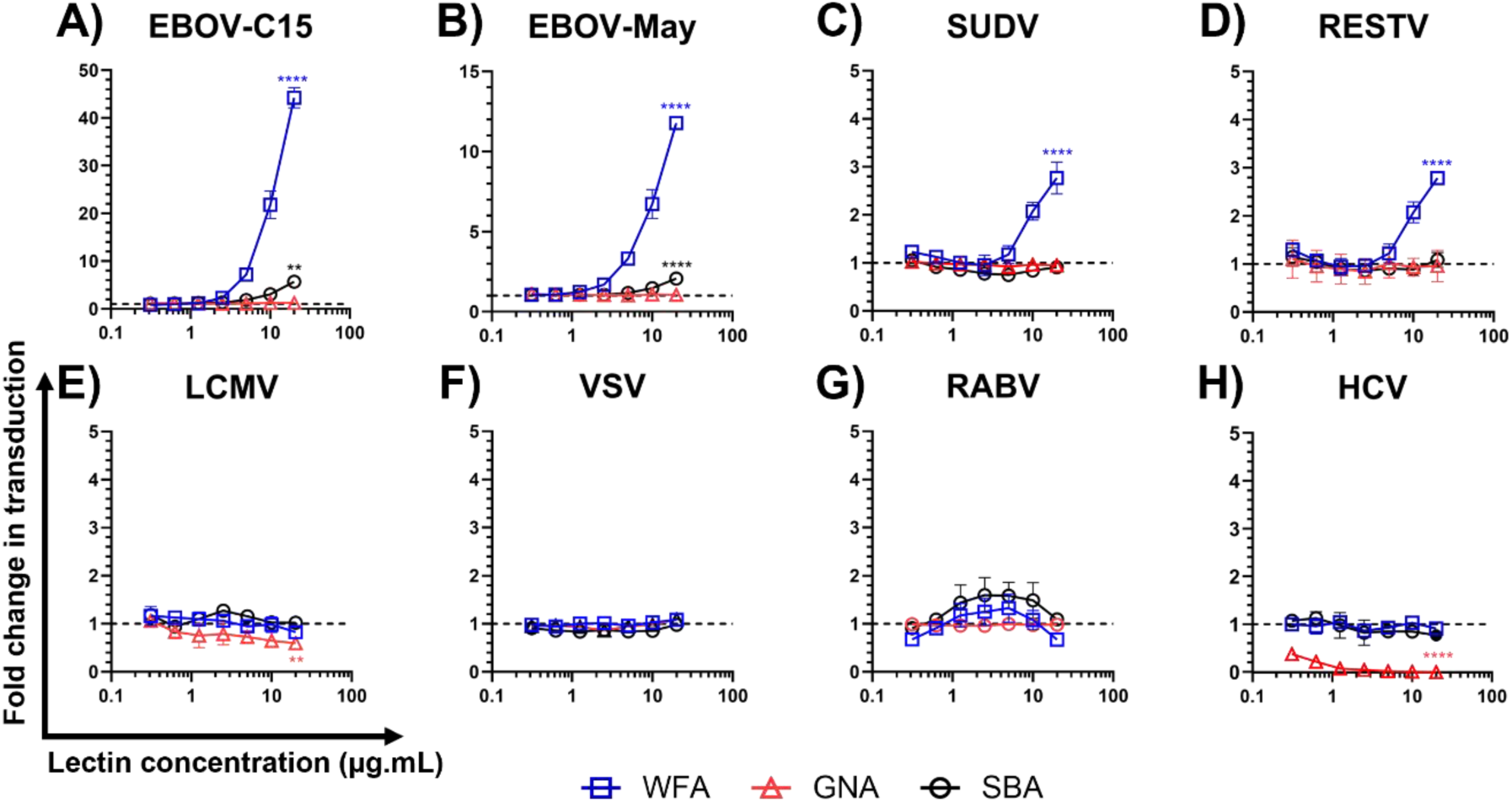
Ebola virus GP-pseudotyped virus transduction is specifically enhanced in the presence of *Wisteria floribunda* agglutinin. Pseudotyped virus particles possessing the envelope glycoproteins of different viral species (including two strains of *Zaire ebolavirus*) were incubated with WFA, GNA or SBA prior to infection of human hepatoma (HuH7) cells. Each lectin was tested in a two-fold dilution series in triplicate with a starting concentration of 20 µg.mL^-1^ . Data are reported as fold change in transduction relative to a no-lectin control group. EBOV-C15; Makona *Zaire ebolavirus*, EBOV-May; Mayinga *Zaire ebolavirus*, SUDV; *Sudan ebolavirus*, RESTV; *Reston ebolavirus*, LCMV; Lymphocytic choriomeningitis virus, VSV; *Indiana vesiculovirus,* RABV; Rabies virus, HCV; hepatitis C virus. The y-axis differs between graphs to aid visualisation. Statistical significance between the effect observed at the highest concentration and a no lectin control was determined by a one-way ANOVA followed by Dunnett’s multiple comparison test, ** p<0.01, **** p<0.0001.

All pseudotypes were able to transduce HuH7 cells (Supplementary figure 2A). Incubation of filovirus PVs with WFA prior to addition to target cells resulted in a dose-dependent increase in viral entry (Figure 1A-D). At a WFA concentration of 20 µg.mL^−1^, the *Zaire ebolavirus* GP PVs displayed a significant mean fold increase of 44.2 (p<0.0001) and 11.8 (p<0.0001) for EBOV-C15 and EBOV-May, respectively (Figure 1A-B). SUDV and RESTV also displayed enhanced viral entry when compared to a no lectin control group (Figure 1C-D). These PVs showed enhancement at a reduced level compared to the EBOV PVs, with a mean fold increase in transduction of 2.8 and 2.9, respectively. No change in entry was observed for any other virus in response to the presence of WFA (Figure 1E-H).

Consistent with previous reports, GNA neutralized HCV PV entry^43,44^ at all concentrations tested (Figure 1H). At a concentration of 20 µg.mL^−1^, GNA exhibited a 0.4-fold reduction in LCMV PV entry which was significant compared to a no lectin control (p<0.01) (Supplementary figure 2B). No significant effects of GNA were observed toward any other PVs in the panel. Following treatment with SBA lectin at a concentration of 20 µg.mL^−1^, HCV PVs displayed a mean 0.23 fold reduction in transduction (p<0.05, Supplementary figure 2B) which contrasted to EBOV-C15 and EBOV-May PVs, showing fold increases of 4.7 and 1.1, respectively. The presence of SBA did not impact the entry of other PVs in the panel.

### Lectin enhancement is mediated by direct interaction with viral glycoproteins

To define how WFA enhancement of EBOV entry occurs, EBOV PV entry assays were performed with HuH7 cells that were treated with lectins either before, during, or after infection with pseudotypes (Figure 2A-B). Enhancement of both EBOV-C15 and EBOV-May entry was only observed when cells were co-treated with both PVs and lectin (Figure 2A-B). The mean fold increase in transduction was found to be statistically significant for both EBOV-C15 and EBOV-May when compared to the untreated control group (p<0.01 and p<0.001, respectively, Figure 2A-B). To confirm that binding of WFA to EBOV pseudotypes occurred directly, a lectin-capture ELISA was performed in which pelleted PV samples were incubated on an ELISA plate coated with WFA and detected using the KZ52 antibody. WFA captured both EBOV-C15 and EBOV-May PV particles (Figure 2C). The specificity of this interaction was demonstrated by performing the enhancement of entry experiment with EBOV-C15 PVs in the presence of increasing concentrations of saccharides (GalNAc, GlcNAc, mannose), which are specific ligands for different lectins. Enhancement was reduced as increasing concentrations of GalNAc were incubated with the WFA, with no effect observed when using either GlcNAc or mannose (Figure 2D). No effect on transduction was observed with the highest concentration of saccharide alone (Figure 2D). No effect of the sugars on entry of VSV PVs was observed (Supplementary figure 3).

**Figure 2.**
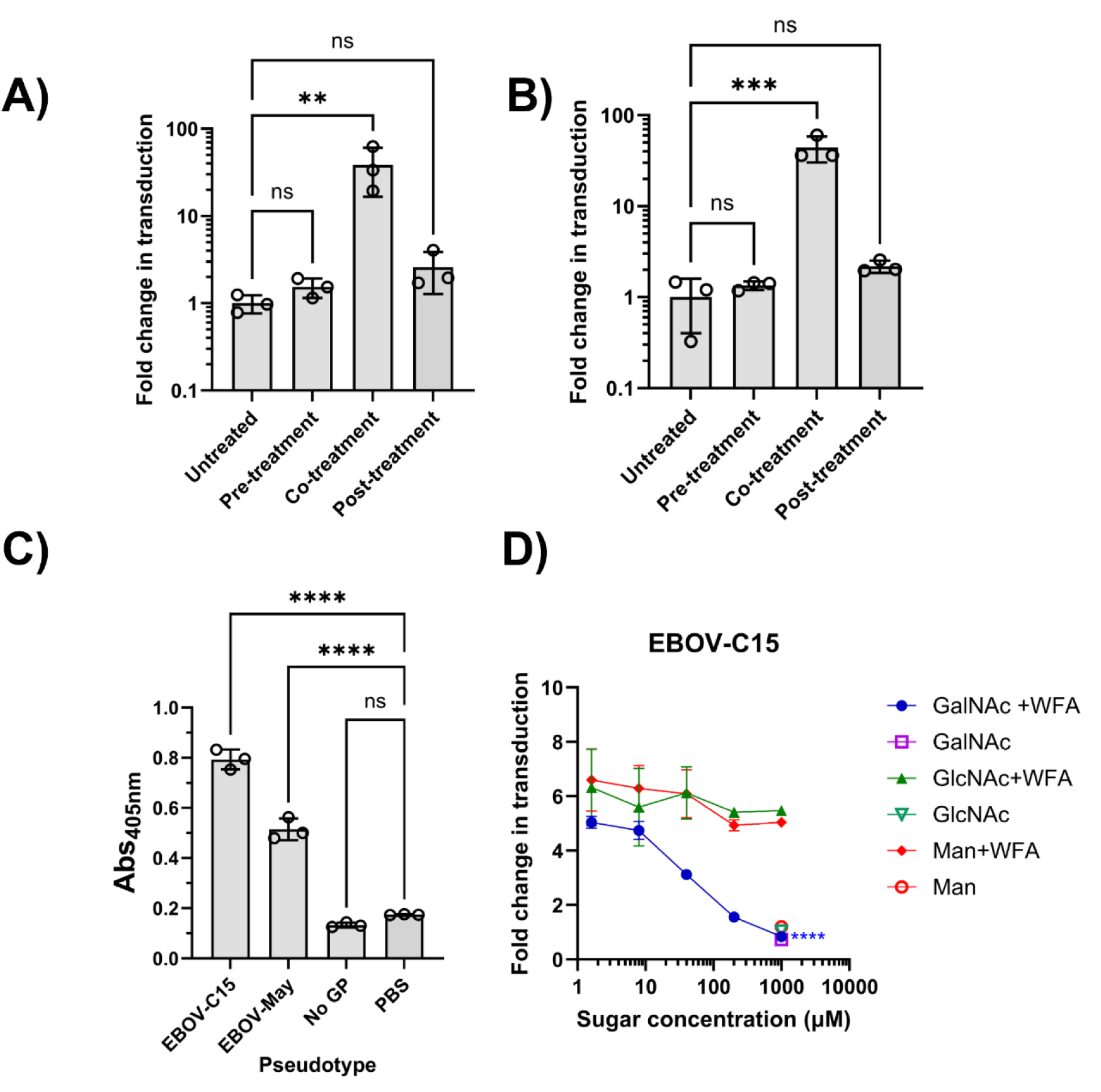
Direct interaction of WFA and EBOV GP results in enhancement. WFA was added at a final concentration of 10 µg.mL^−1^ to HuH7 cells at different stages of EBOV-C15 (**A**) and EBOV-May (**B**) PV entry. Cells were pre-treated with WFA prior to addition of EBOV PVs. In the co-treatment condition, EBOV PVs and WFA were added simultaneously and incubated for 1 hour. Post-treated WFA was added post EBOV PV incubation. PV entry was measured by the luciferase reporter expression and RLU values were compared with the untreated group. **C)** Binding of pseudotypes to WFA was assessed by ELISA. Virus particles were incubated with coated WFA and binding revealed with incubation with anti-GP antibodies. Particles lacking glycoprotein were used as a negative control. **D)** Specificity of interaction was determined by incubating GalNAc, GlcNAc or mannose with WFA during the co-treatment with EBOV-C15 pseudotype entry into HuH7 cells. Statistical significance was determined using one-way ANOVA followed by Dunnett’s multiple comparison test, p<0.01 (**), p<0.001(***), p<0.0001(****), p>0.05 (ns).

### Enhancement of EBOV entry by WFA is species independent but NPC1-dependent

To determine if the enhancing effect of WFA upon EBOV PV entry was specific to human cells, further entry assays were performed with the EBOV-permissive HypLu/45.1 cell line derived from foetal lung tissue from the bat species *Hypsignathus monstrosus*^45^ (a potential reservoir of EBOV^46^). In agreement with the observed effect of WFA on EBOV-C15 entry in HuH7 cells (Figure 3A), entry of EBOV-C15 into HypLu/45.1 cells was enhanced by the presence of WFA, albeit with reduced effect (Figure 3B). We also investigated whether WFA-mediated enhancement of EBOV PV entry was independent of the normal NPC1-dependent entry pathway. A previously validated NPC1 knockout U2OS cell line was utilised^7^. The enhancing effect of WFA on EBOV-C15 entry in U2OS cells was comparable to that observed for HuH7 cells (Figure 3C). U2OS-NPC1 KO cells did not permit EBOV-C15 PV entry (Figure 3D). WFA was unable to facilitate EBOV PV entry into these cells, demonstrating that the process supporting entry facilitated by WFA remained NPC1-dependent. Unexpectedly, for the control VSV PVs, the inclusion of WFA during infection was found to enhance infection when using U2OS-NPC1-KO cells as targets (Figure 3D). This contrasted with all previous experiments conducted with both human and bat cell lines.

**Figure 3.**
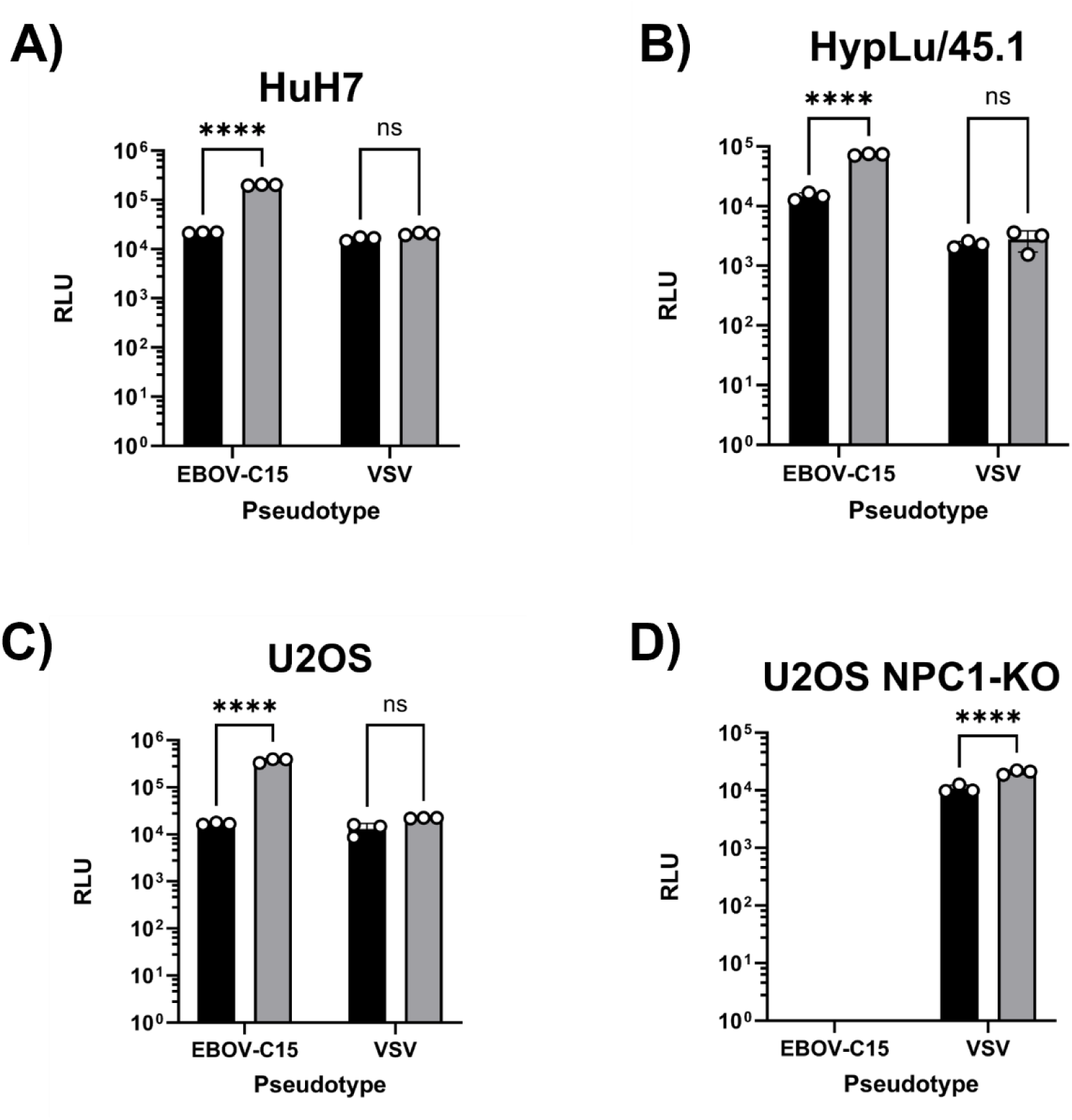
WFA-mediated enhancement of entry occurs in different species and is NPC1-dependent. Four mammalian cell lines, HuH7 (human, **A**) and HpyLu/45.1 (bat, **B**), human osteosarcoma cells (U2OS, **C**) and a validated U2OS NPC1-knock-out cell line (**D**) were transduced with pseudotypes possessing the glycoproteins of Ebola C15 or VSV, +/-WFA at a concentration of 10 µg.mL^−1^. Black bars indicate the control treated pseudotypes and grey bars indicate the WFA-treated pseudotypes. Significance was determined by one-way ANOVA followed by Sidak’s multiple comparison test, ns p>0.05, **** p<0.0001.

### WFA-mediated enhancement of EBOV entry is modulated by the presence of individual polymorphic *N*-linked glycosylation in the glycan cap

The Zaire EBOV-GP undergoes extensive glycosylation, with 17 *N*-linked glycan sites across the protein. To identify key glycosylation sites that mediate WFA enhancement of EBOV PV entry, transduction assays were performed using five previously described naturally-occurring variants of the EBOV-C15 GP that lack *N-*linked glycosylation sites^47^ (Figure 4). These variants all possess an A82V mutation, located in the head region of GP_1_, that became established early during the 2013-16 EBOV epidemic^48^ and is thought to enhance viral membrane fusion^49,50^. This variant was labelled ‘B1’. In addition, variant B12 encodes a T206M mutation while variants B13, B14 and B16 each possess a T230A mutation; both of which disrupt *N*-linked glycosylation sequons, preventing glycosylation at residues N^228^ and N^204^, respectively (Supplementary table 1). The B1 variant was included as a control for the A82V mutation, as this variant was not predicted to have an altered *N*-linked glycosylation pattern compared to the EBOV-C15 control, and preliminary experiments showed that B1 PVs exhibited the same pattern of enhancement as EBOV-C15 (Supplementary figure 4). In line with previous observations, each of the variant EBOV PVs exhibited enhanced entry in the presence of WFA (Supplementary figure 5). When compared to a no lectin control group, B1 and B12 showed a significant fold change of 23.8 (p<0.01) and 22.5 (p<0.01), respectively (Figure 4A). Variants lacking glycosylation at residue N^228^ - B13, B14, and B16 – exhibited lower levels of enhancement, with mean fold changes of 12.3 (p<0.01), 14.2 (p<0.05) and 14.5 (p<0.05), respectively (Figure 4A; Supplementary figure 5). Surprisingly, each variant was also significantly enhanced by the SBA lectin at a concentration of 10 µg.mL^−1^, and GNA lectin did not impact EBOV or VSV PV entry (Supplementary figure 5).

**Figure 4.**
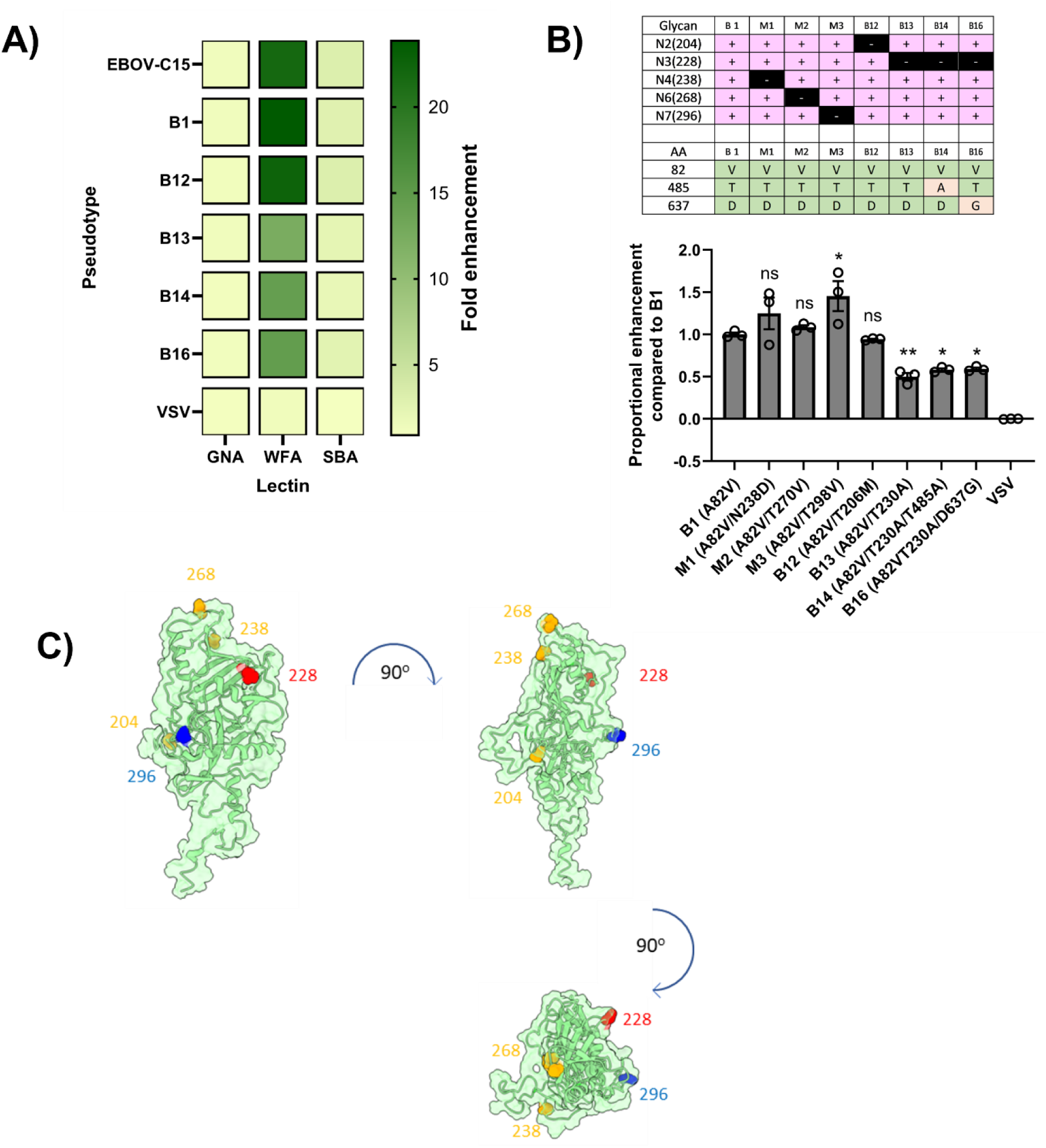
WFA-mediated enhancement of entry is modified in naturally occurring Ebolavirus variants. **A)** Infection assays using HuH7 cells were performed using GPs from naturally occurring EBOV isolates with predicted altered *N*-linked glycosylation sites from the wildtype C15. Fold change between each lectin was compared to a no lectin control. **B)** Three *in vitro*-generated glycan mutants M1, M2, M3 were compared with variants B1, B12, B13, B14 and B16 in the enhancement assay at 10 µg.mL^−1^. Differences were determined using one-way ANOVA and a Dunnett’s multiple comparison test, ns p>0.05, * p<0.05, ** p<0.01. **C)** Representation of the sites of *N*-linked glycosylation on a model of the Mayinga EBOV GP_1_ protein (produced using I-TASSER using the Makona C15 amino acid sequence modelled onto PDB structure 6VKM). Sites where removal of the glycan did not affect enhancement are highlighted in yellow, one (296) that increased enhancement in blue, and one (228) that reduced enhancement highlighted in red.

To quantify the effect of polymorphisms in the EBOV GP_1_, these five naturally occurring variants were analysed alongside three previously-described *in vitro*-generated mutants^51^ made in a B1 background, M1 (N238D), M2 (T270V) and M3 (T298V), resulting in loss of glycosylation at N^238^, N^268^ and N^296^, respectively. Normalising for the differences in transduction observed for each mutant, enhancement was compared to the B1 reference (Figure 4B). All pseudotypes possessing the T230A polymorphism had significantly reduced enhancement (B13, p<0.01; B14, p<0.05; B16, p<0.05), while entry enhancement of the T298V mutant was significantly increased (p<0.05) (Figure 4B). When modelled on a structure of the GP_1_ molecule based on PDB accession number 6VKM, the residues that impacted on WFA-mediated enhancement were found to be separated from those that did not (Figure 4C). When visualised with the NPC1 binding pocket, the distance between the N^228^ and the I^113^ within the NPC1 binding site was measured to be ∼18Å.

In addition to individual amino acid variants, a modified clone of EBOV-C15 was generated possessing a complete MLD deletion (EBOV-C15-ΔMLD, removing aa305-485 of GP_1_). This contained eight fewer *N*-linked glycosylation sites (Supplementary table 1), and removed the majority of O-linked glycosylation sites^13^. In line with previous reports^9^ the EBOV-C15-ΔMLD mutant displayed a greater level of infectivity compared to the wild-type EBOV-C15 PVs (Supplementary figure 6). Both EBOV-C15 and EBOV-C15-ΔMLD PVs were susceptible to enhancement by WFA lectin (Figure 5A-B). Interestingly, at a WFA concentration of 10 µg.mL^− 1^ the removal of the MLD significantly increased the enhancing effect of WFA from a mean fold change of 8.74 to 12.62 (p<0.0001, Figure 5C). Additional enhancement by SBA was also revealed in the absence of the MLD from a mean fold change in transductions of 3.07 to 5.92 (p<0.0001) (Figure 5C).

**Figure 5.**
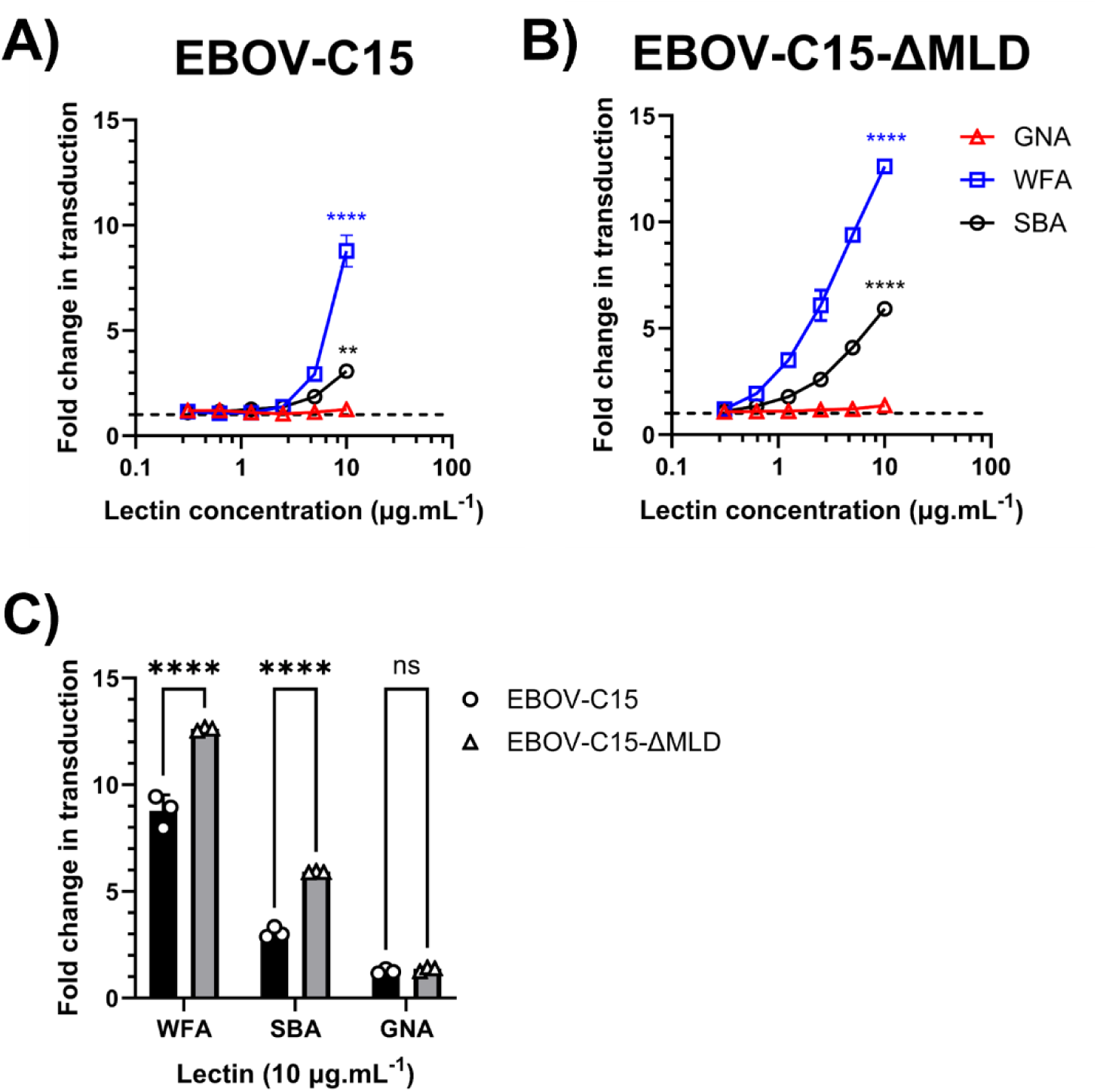
Enhanced entry of mucin domain deleted mutant of Ebolavirus C15 by WFA. Transduction assays with HuH7 cells were performed in triplicate with EBOV-C15 (**A)** and EBOV-C15-ΔMLD (**B**) PVs using two-fold serially diluted lectins starting from 10 µg.mL^−1^. **C)** Fold change in transduction of EBOV-C15 and EBOV-C15-ΔMLD at a lectin concentration of 10 µg.mL^−1^. Black bars: pseudotypes possessing the EBOV-C15 GP; grey bars: pseudotypes possessing the EBOV-C15-ΔMLD GP. Comparisons were determined by one-way ANOVA followed by Sidak’s multiple comparison test, p> 0.05 (ns), p<0.0001 (****).

### Reduced antibody-mediated neutralization of WFA-treated EBOV PVs

To determine if the binding of WFA to the EBOV GP complex resulted in altered GP antigenicity, the impact of WFA on antibody-mediated neutralisation and binding was investigated. Neutralisation assays were performed with the well-characterised conformation-dependent anti-GP nAb, KZ52^6^ against EBOV PVs in the presence of WFA. EBOV PVs bearing C15-GP or ΔMLD-GP were susceptible to neutralisation by KZ52 in both the presence and absence of WFA (Figure 6A). For EBOV-C15 PV, the IC_50_ value of KZ52 was 5.85 µg.mL^−1^ in the presence of WFA, and 3.86 µg.mL^−1^ without WFA (Figure 6A, Supplementary table 2). The IC_50_ values of KZ52 for ΔMLD-GP PV were 16.76 µg.mL^−1^ and 2.07 µg.mL^−1^ in presence or absence of WFA, respectively (Figure 6A, Supplementary table 2). Modelling of the interaction of WFA with contacts at the N^228^ residue in the context of a trimeric presentation of the trimer of GP_1,2_ heterodimers in complex with KZ52 Fabs (Figure 6B) highlighted that, in the context of full GP_1,2_, binding of WFA to the EBOV spike protein may occur without directly blockading the KZ52 epitope. However, certain orientations of the tetrameric lectin could potentially compete with the antibody for binding.

**Figure 6.**
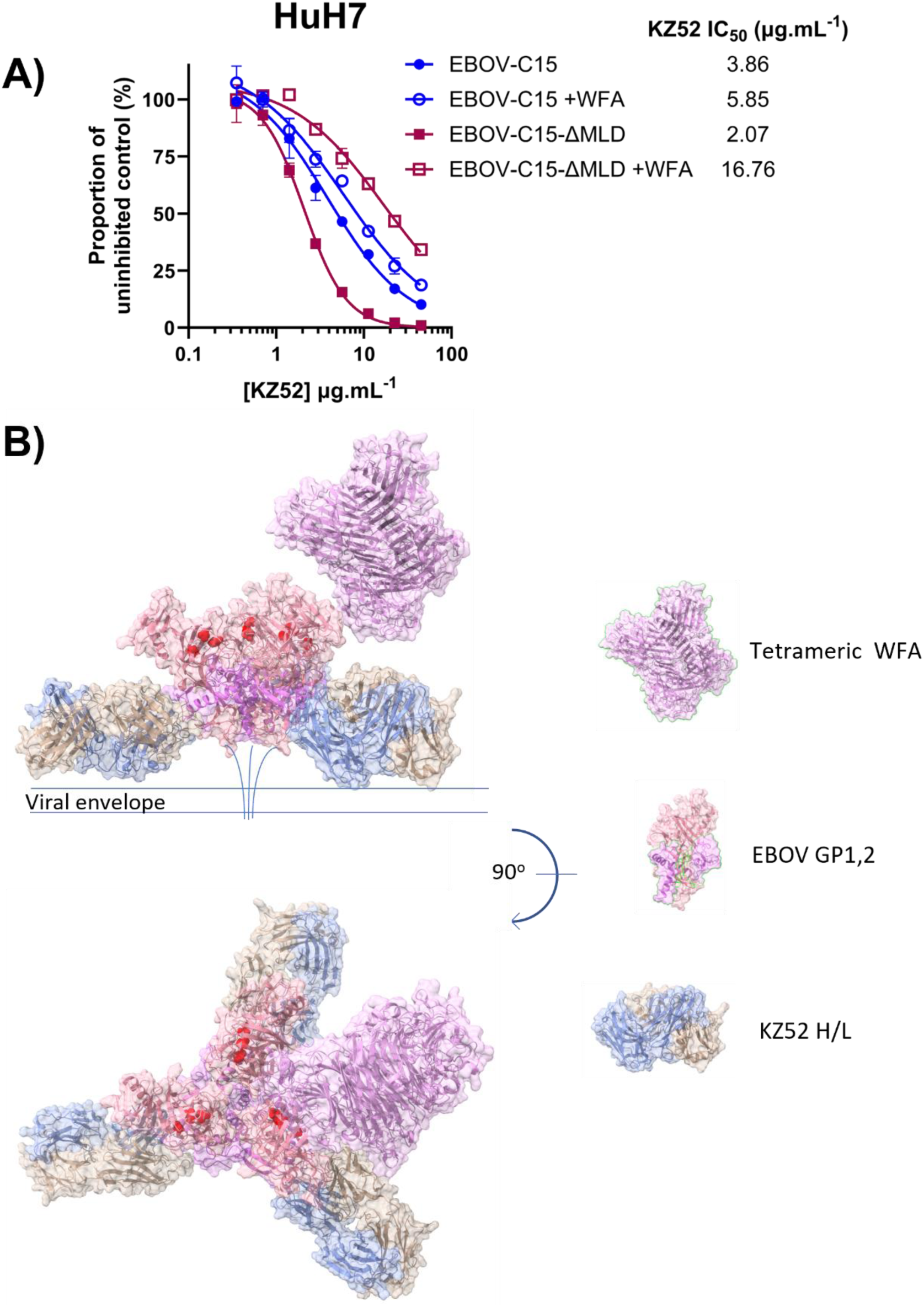
Modelling of WFA binding to EBOV GP. **A)** mAb KZ52 neutralisation of EBOV-C15 pseudotype entry into HuH7 cells in the presence or absence of WFA lectin. EBOV PVs possessing the C15 or C15-ΔMLD GPs were incubated with a two-fold serial dilution of KZ52 in the absence or presence of WFA, at a final concentration of 10 µg.mL^−1^. Data was normalised to a no antibody control. **B**) Modelling of the trimeric EBOV-GP in complex with KZ52 (PDB 3CSY) with one possible configuration of the tetrameric WFA molecule (PDB 5KXB) in close association with N^228^ of GP_1_ was performed using ChimeraX. Amino acid residues implicated in the NPC1 interaction are highlighted in red.

## Discussion

This study provides the first evidence of plant-derived lectins enhancing EBOV-GP mediated cell entry. Initial testing of three different plant lectins identified WFA as a potent enhancer of filovirus GP-mediated cellular entry. Two lectins were observed to enhance infection (WFA and SBA), but because the effect was most potent for WFA this was investigated in more detail. However, the GalNAc- or Gal-specific SBA may exhibit similar properties to WFA. It remains to be determined why some plant lectins, such as mannose specific BanLec^37^, inhibit infection, but others have enhancing effects on filovirus entry. Of the filovirus species tested, EBOV displayed the greatest levels of enhancement in the presence of WFA, suggesting amino acid sequence-dependent activity. Enhancement was also observed in naturally occurring variants of the EBOV-GP that display altered patterns of *N*-linked glycosylation. Furthermore, greater WFA-mediated enhancement occurred following the deletion of the MLD region, which contrasts with the MLD-dependent enhancement previously described for human ficolin-1^32^. Together this indicates that the binding of WFA to EBOV-GP occurs at GalNAc-containing glycans present in the glycan cap of GP_1_. The glycosylation of GP_1_ is highly heterogeneous, with many complex glycans associated with each site. In human cells, each *N*-linked glycan site is predicted to possess, on average, 12 glycan compositions^13^. The data presented here supports the conclusion that carbohydrate structures possessing accessible LacdiNAc or GalNAc are present on the surface of the GP_1_ glycan cap. Given the restricted range of sugars attached to N^257^ in GP ^13^, and the impact of variants observed here, it is most likely that these sugars are present at positions N^40^, or N^228^, or to *O*-linked sugars present in the glycan cap^52^. The confirmation that removing N^228^ reduced, but did not ablate enhancement confirms that this is a complex effect mediated by multiple glycans. While N^40^ is completely conserved across all filovirus species, N^228^ is present in EBOV and RESTV, but not SUDV GP (Supplementary figure 1). The reduction in enhancement observed for the N^228^ mutant suggests that this residue contains a glycan presenting a ligand for WFA. The enhancement of infection for this variant was comparable to that of *Sudan ebolavirus* and *Reston ebolavirus*. Other sites contribute to the enhanced infection, and while direct protein-protein interactions cannot be excluded it is likely that patterns of other conserved glycans contribute to the observed effect. While we did not explore the effect of altering the potential *O*-linked glycosylation sites in this study, it is plausible that these contribute to the enhancing effect of WFA. GalNac-containing *O*-linked sugars have been identified on the GP_1_ by mass spectrometry^52^. It is challenging to accurately predict the number of *O*-linked glycans attached to the GP_1,2_ structure^52^, but two *O*-linked sites in GP_1_ have been well characterised^13^. Further investigations are required to determine the contribution of *O*-linked glycosylation to lectin-mediated entry enhancement.

A potential limitation to this study was the reliance of a retroviral pseudoviral model of EBOV entry. This is a tractable approach to model entry of pathogenic viruses under Containment Level 2 laboratory settings. PVs have been used to model EBOV GP function and processing, and have the benefit of targeting the entry cascade alone. However, there are noted differences between EBOV PVs and authentic virus. For example, EBOV PVs have been shown to be more resistant to the action of EBOV neutralising mAbs compared to live virus^53,54^. Alternative models for studying EBOV, such as a virus-like particles^55^ or VSV-based pseudotypes^53^ could also be used for these studies, which may reproduce the morphology of the virus particles more accurately. However, the glycosylation of EBOV pseudotypes is similar to authentic GP^56^, and these have been used previously to investigate the EBOV entry pathway^57^. The results presented here provide evidence that could be further explored with both a VLP infection assay and live virus cultures to confirm the observed enhancing effect in other models of EBOV entry. The choice of target cells for infection assays may also influence the outcome of experiments. In the present study the HuH7 cell lines utilised for the majority of experiments as they do not express DC-SIGN/L-SIGN, hMGL or LSECtin, but do express high quantities of ASGPR1^58,59^. Our confirmation of the same enhancing effect in U2OS cells, which express much lower levels of ASGPR1, and non-human cells demonstrate that the enhancing effect is likely to be generalisable to all cell types. Further studies are required to assess the enhancing effect of WFA on cells expressing the complete range of membrane-associated lectins that may interact with the EBOV GP.

The observation that WFA-treated EBOV PVs were less well neutralised by the KZ52 mAb indicated that WFA may either partially occlude the KZ52 epitope, or that a change occurs upon WFA binding that alters the overall conformation of the glycoprotein complex, either to impact antibody binding or accelerate entry kinetics. KZ52 recognizes residues 505–514 and 549–556 of GP_2_ and residues 42–43 of GP ^6^. Our modelling of the interaction of the WFA with glycosylation sites in GP demonstrated that binding to the oligomeric GP complex is possible without directly blockading the KZ52 binding site (Figure 6B). It is plausible that conformational changes induced by lectin binding indirectly alter the epitope of this conformation-dependent antibody, and given that the N^228^ residue is located at a site between the NPC1 binding site and the GP_2_ subunit, but not directly overlapping, the blockade by WFA may not be direct. Further structural studies are required to resolve these possible interpretations of our data.

Collectively, the data presented here provide evidence that interactions between GalNAc-specific lectins and the EBOV GP results in changes to the envelope glycoprotein that result in greater binding and fusion through an NPC1-dependent process. It is plausible that enhancement caused by lectins expressed in mammalian cells exhibit a similar mode of action. We propose a model of entry where initial, generalisable interactions with lectins (whether soluble, or membrane-associated) trigger conformational changes that facilitate internalisation and subsequent interaction with NPC1, negating the requirement for proposed lectin receptors^32^ expressed on the surface of cells to mediate virus entry. This is consistent with previous assessment of the role of DC-SIGN in entry^28,60^. The demonstration that the enhancing effect of WFA is more pronounced when using a ΔMLD construct highlights the possibility that lectin interactions may be promoted after the initial cell surface interactions and subsequent cathepsin B/L cleavage of GP_1_ prior to NPC1 interaction.

## Material and Methods

### Plasmids for virus glycoprotein expression

A codon optimised sequence for the Makona isolate, Kissidougou-C15 GP (Accession AHX24649.2) was synthesised and inserted into a pcDNA3.1 (Invitrogen). The generation of variant GP sequences from Makona C15 by *in vitro* mutagenesis was described in a previous study^47^. The plasmids encoding Ebola Mayinga GP and other *Ebolavirus* species were kind gifts from G. Kobinger (Galveston National Laboratory, University of Texas). Hepatitis C virus H77 E1/E2 construct has been previously described^61^. The RABV-G protein construct was a kind gift of Dr E. Wright (University of Sussex), and the LCMV-G protein-encoding plasmid was generated in house. The pM2D plasmid encoding the VSV-G glycoprotein was obtained from AddGene (plasmid # 12259). The C15-ΔMLD construct was created by inverse PCR of the EBOV-C15 GP plasmid (primers available upon request) to generate a deletion of amino acids 305-485. Individual amino acid point mutants were generated using a Q5-SDM kit (NEB) using the B1 variant as template.

### Production of pseudotype virus (PV) particles

Retroviral pseudotype particles were produced as descried previously^47^. In brief, 1.5 million HEK-293T cells were seeded in a Primaria coated 10 cm dish (Corning) and incubated overnight. Dishes were transfected with 2 µg of MLV-gag/pol encoding packaging plasmid phCMV-5349, 2 µg of the MLV-Luc plasmid pTG126 and 2 µg of plasmid encoding the viral envelope protein of interest. For generation of EBOV-PPs, cells were transfected with 0.2 µg of plasmid encoding EBOV-GP. Negative control ‘No GP’ particles were generated by transfection without a viral envelope protein encoding plasmid. For each transfection, plasmids were diluted in serum-free Opti-MEM (Gibco), mixed with 24 µL PEI (Polysciences) in a final volume of 600 µL and incubated for 1 h at room temperature. Cell media was replaced with 7 mL Opti-MEM and plasmid-PEI solution was added in a dropwise manner. Following a 6 h incubation at 37 °C, the Opti-MEM was replaced with 10 mL DMEM. Cell supernatants were harvested 72 h post transfection and passed through a 0.2 µM syringe filter (Sartorius) and stored at 4 °C.

### Transduction assays

HuH7 cells^62^ were used for the majority of infection experiments. For specific infection in NPC1-KO cells, previously described U2OS cell lines were used for infection. These cells were seeded at a density of 1.5 ×10^4^ cells/well in a 96 well white plate (Corning) and grown overnight. Media was aspirated from the wells and replaced with 100 µL of PV-containing supernatant. For incubation with lectin, WFA (Sigma L1516), SBA (Sigma L1395) or GNA (Sigma L8275) were diluted from a 10 mg.mL^−1^ stock solution to the required final concentration in the pseudotype-containing media. Incubation with lectins were performed for 1 hour before infecting target cells. The inoculum was removed after 4 h and 200 µL of DMEM was added per well and cells incubated for 72 h. To measure luciferase activity, media was removed, and cells were lysed in 50 µL cell lysis buffer (Promega) and incubated at room temperature for 30 min with gentle agitation. Plates were then read using a BMG Fluostar Omega plate reader, primed with Luciferase substrate with gain set to 3600.

### Antibody Neutralization Assays

The KZ52 antibody was produced using a HEK-293 Freestyle expression system (Thermo Fisher Scientific). In brief, plasmids encoding the KZ52 heavy chain and light chain (pORIgHIB and pORIgLB^63^, respectively) were mixed with PEI at a ratio of 1:3 and incubated at room temperature for 1 h. Cell density was adjusted to 1 ×10^6^ cells.mL^−1^ and plasmid-PEI mix was added. Transfected cultures were incubated for 72 h and media was harvested. KZ52 concentration was determined using an in-house IgG ELISA assay using anti-light chain antibody to capture the IgG and an anti-heavy chain HRP-conjugated antibody for detection. KZ52 was incubated with EBOV PVs with or without WFA for 1 h at room temperature and 100 µL was added per well to HuH7 cells. Cells were incubated with EBOV PVs for 4 h at 37°C and media was replaced with fresh DMEM. Cells were grown for 72 h and transduction was measured using the previously described luciferase assay.

### ELISA Binding Assays

Pseudotypes were purified from transfected HEK 293T cell supernatants by ultracentrifugation at 107,000 *x* g for 180 minutes through a 20% sucrose cushion. Pellets were resuspended and analysed by western blot using an anti-MLV p30 antibody (Sigma). Following quantification, equal quantities of pseudotypes were used for ELISA, either directly coated to NUNC Maxisorp microtitre plates or by capturing on plates coated at 10 µg.mL^−1^ with WFA. Pseudotypes were then detected using mAb KZ52, either in the presence or absence of WFA, incubated concurrently. Detection of bound KZ52 was achieved using an anti-human IgG alkaline phosphatase conjugated antibody (Sigma, A3187), followed by development with pNPP (Sigma, N1891) and reading at 405 nm.

### Statistics

Data was analysed using GraphPad Prism software (version 10.3.1). For determining the IC_50_ values of KZ52, data were normalised to a no antibody control group and nonlinear regression was performed using the log(agonist) vs. response-variable slope (four parameters model, with the bottom parameter constrained to a value of 0). Multiple comparisons were performed with a one-way ANOVA, using either Sidak’s test, or Dunnett’s test to account for multiple comparisons where appropriate. Where only two datasets were compared, a Student’s t-test was used.

### Structural modelling

For identifying *N*-linked glycan sites, the amino acid sequence of the EBOV C15 GP1 protein was modelled using I-TASSER (https://zhanggroup.org/I-TASSER/) using PDB structure 6VKM as a scaffold. Representation of the generated model was visualised in ChimeraX (https://www.cgl.ucsf.edu/chimerax/). Modelling of the oligomeric structure of the trimeric GP_1,2_ in complex with mAb KZ52 (PDB 3CSY) and the tetrameric WFA (5KXB) proteins was performed with ChimeraX.

## Acknowledgments

We are grateful for the provision of glycoprotein-expression constructs from the laboratories of Edward Wright and Gary Kobinger. We are also grateful to Kartik Chandran and colleagues for the provision of the U2OS NPC1-knockout cell line. We would also like to acknowledge the contribution of undergraduate research project students who contributed to preliminary experiments for this study. This work was funded by the Medical Research Council UK (grant MR/S009434/1).

---

**Supplementary Table 1.** Positions of *N*-linked glycan sites in the EBOV GP.

| Glycan | Variant |  |  |  |  |  |  |  |  | Location | Cross-species |
| --- | --- | --- | --- | --- | --- | --- | --- | --- | --- | --- | --- |
|  | B1 | M1 | M2 | M3 | B12 | B13 | B14 | B16 | ΔMLD |  |  |
| N1(40) | + | + | + | + | + | + | + | + | + | Base | Y |
| N2(204) | + | + | + | + | - | + | + | + | + |  | Y |
| N3(228) | + | + | + | + | + | - | - | - | + | Cap | N |
| N4(238) | + | - | + | + | + | + | + | + | + | Cap | Y |
| N5(257) | + | + | + | + | + | + | + | + | + | Cap | Y |
| N6(268) | + | + | - | + | + | + | + | + | + | Cap | Y |
| N7(296) | + | + | + | - | + | + | + | + | + | Cap | N |
| N8(317) | + | + | + | + | + | + | + | + | - | MLD | Y |
| N9(333) | + | + | + | + | + | + | + | + | - | MLD | Y |
| N10(346) | + | + | + | + | + | + | + | + | - | MLD | N |
| N11(386) | + | + | + | + | + | + | + | + | - | MLD | N |
| N12(413) | + | + | + | + | + | + | + | + | - | MLD | Y |
| N13(436) | + | + | + | + | + | + | + | + | - | MLD | Y |
| N14(454) | + | + | + | + | + | + | + | + | - | MLD | N |
| N15(462) | + | + | + | + | + | + | + | + | - | MLD | Y |
| N16(563) | + | + | + | + | + | + | + | + | + | GP2 | Y |
| N17(618) | + | + | + | + | + | + | + | + | + | GP2 | Y |

**Supplementary Table 2.** IC_50_ values of KZ52 against EBOV-C15 and EBOV-C15-ΔMLD in the presence and absence of WFA lectin at a concentration of 10 µg.mL^−1^ with 95% CI.

|  | KZ52 |  |
| --- | --- | --- |
|  | IC<sub>50</sub> (μg.mL<sup>-1</sup>) | 95% CI |
| **EBOV-C15** | 3.86 | 2.82 to 4.89 |
| **C15+ WFA** | 5.85 | 4.09 to 7.54 |
| **EBOV-C15-ΔMLD** | 2.07 | 1.86 to 2.28 |
| **EBOV-C15-ΔMLD + WFA** | 16.76 | 13.96 to 19.65 |

**Supplementary Figure 1.**
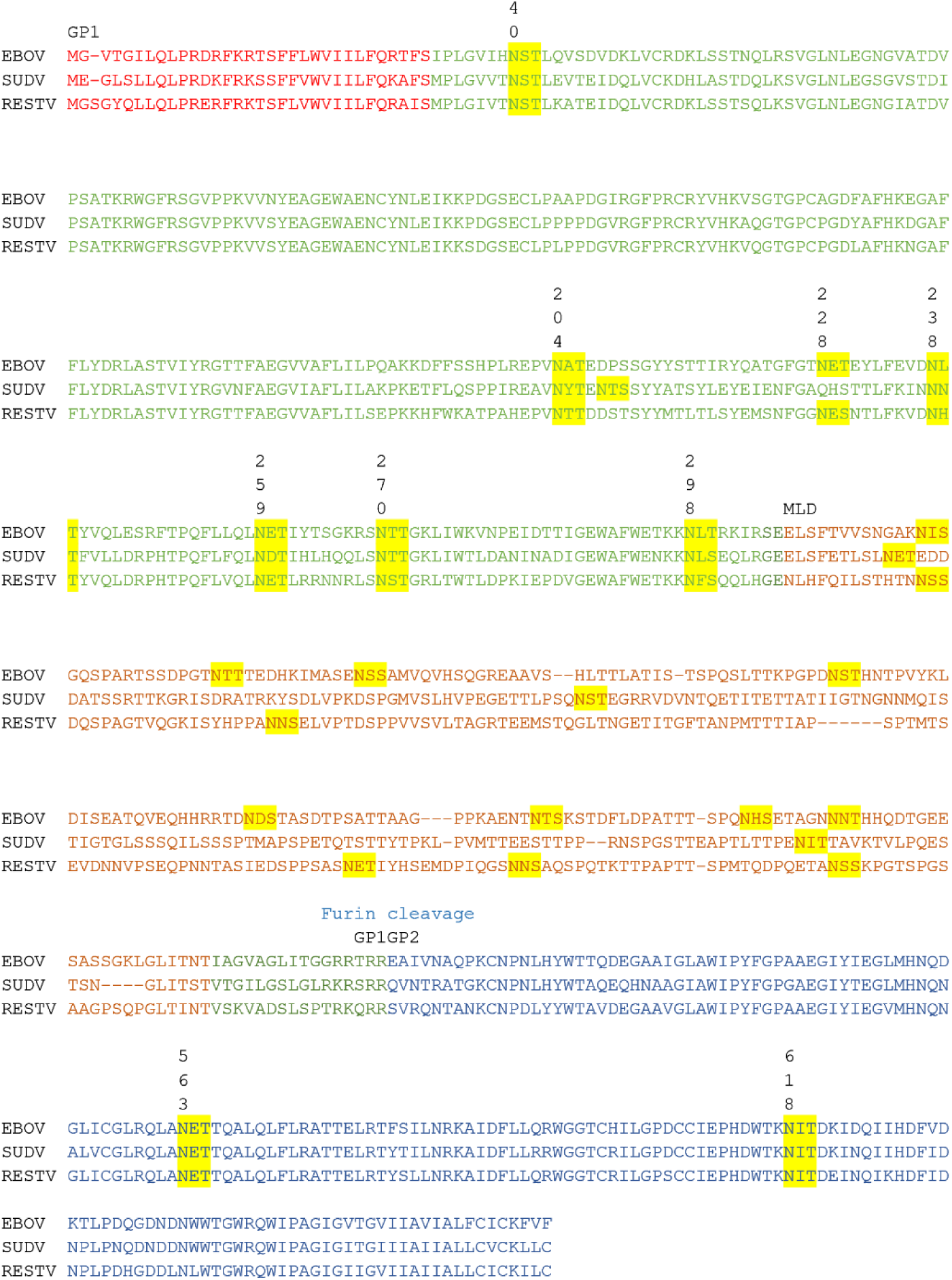
Alignment of *Zaire ebolavirus* (EBOV), *Sudan ebolavirus* (SUDV) and *Reston ebolavirus* (RESTV) GP_1,2_ amino acid sequences. The signal peptide is indicated in red text. The mature GP_1_ is highlighted in green, with the mucin-like domain indicated in brown GP_2_ is highlighted in blue. N-linked glycosylation sequons are highlighted in yellow, with sites in GP_1_ and GP_2_ indicated with the numbering from the EBOV genome sequence (Uniprot accession number Q05320).

**Supplementary Figure 2.**
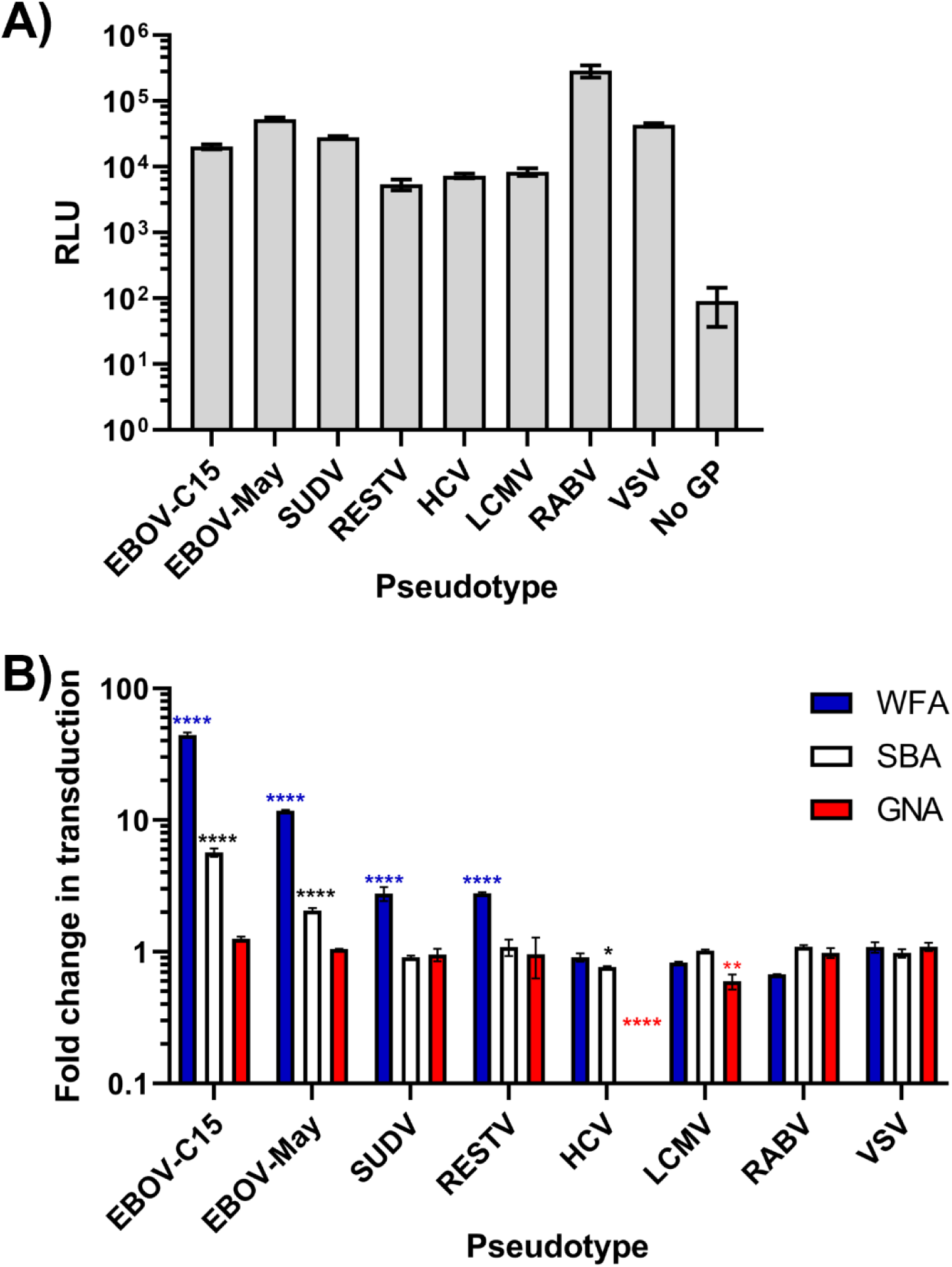
**A)** Luciferase activity of HuH7 cells transduced by the panel of PVs. **B)** Comparison between the mean fold change in transduction of each PV with lectin at a concentration of 20 µg.mL^−1^ and a no lectin control. Statistical significance was determined by one-way ANOVA followed by Dunnett’s multiple comparison test, p<0.05 (*), p<0.01 (**), p>0.0001 (****). Comparisons with p>0.05 are not labelled.

**Supplementary Figure 3.**
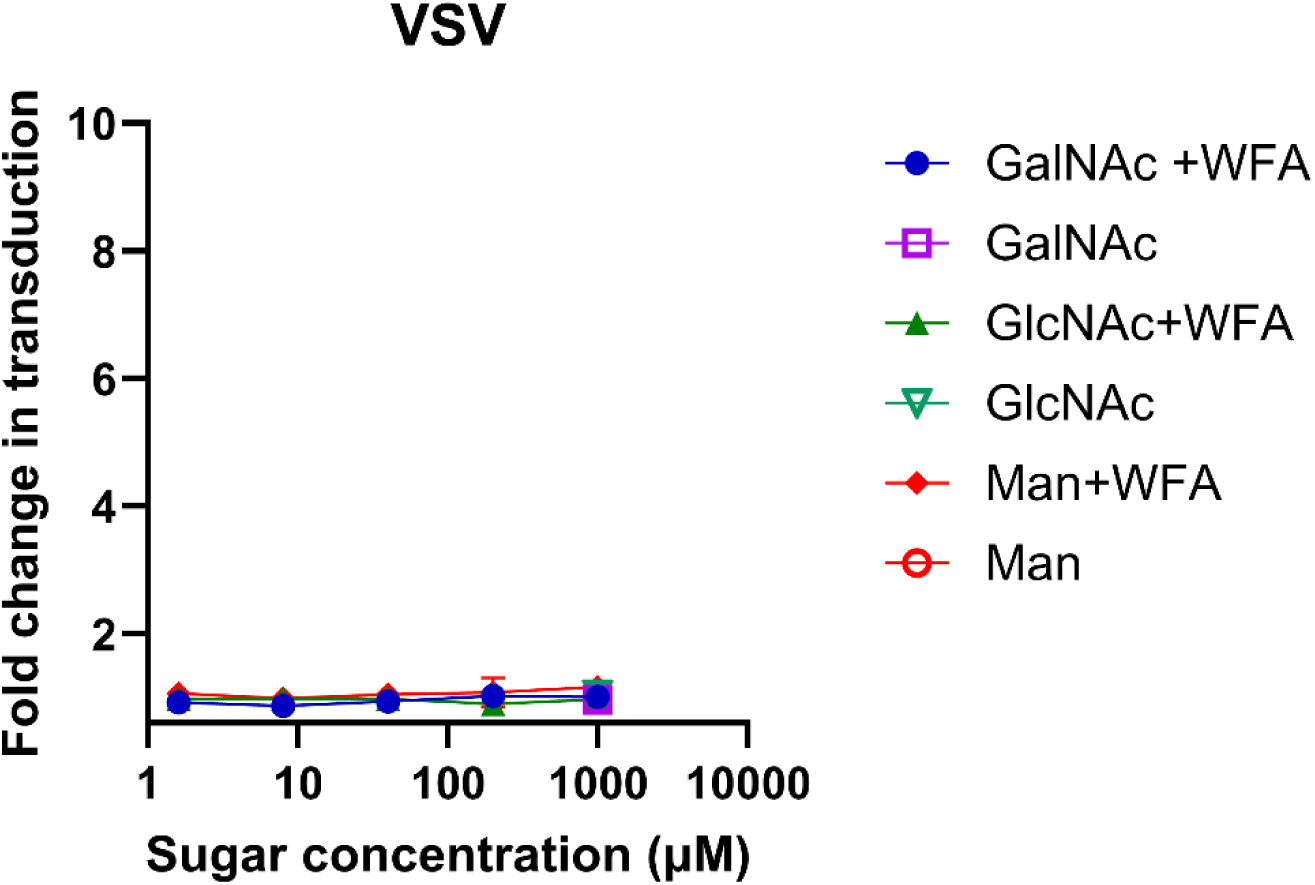
Effect of sugars (GalNAc, GlcNAc, mannose (Man)) on transduction of HuH7 cells with VSV pseudotypes. WFA was used at a concentration of 10µg.mL^-1^ . Statistical significance was determined by one-way ANOVA on samples treated with the greatest concentration of each sugar, followed by Dunnett’s multiple comparison test.

**Supplementary Figure 4.**
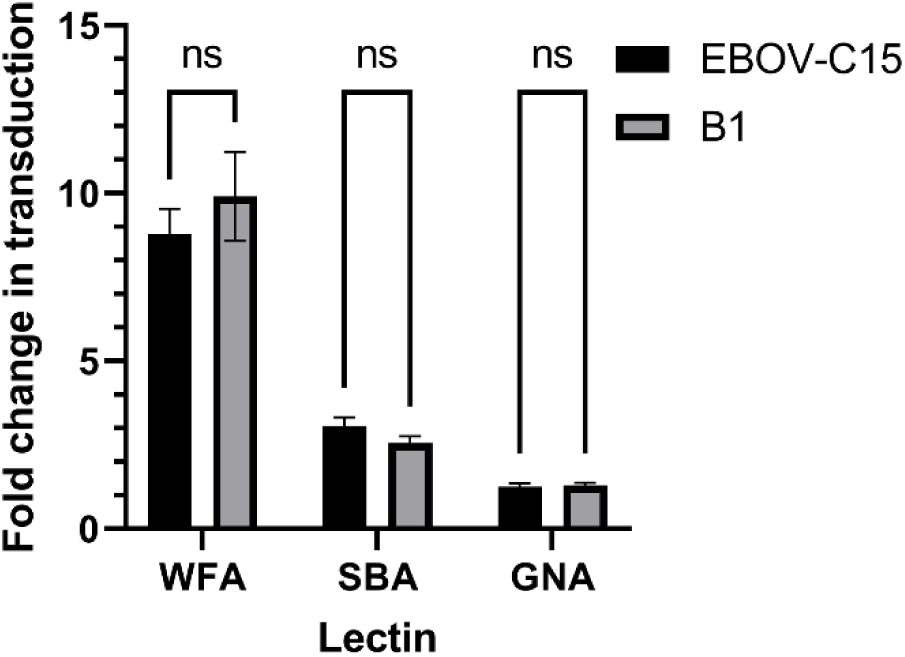
Enhancement of transduction of HuH7 cells with *Ebolavirus* GP variants EBOV-C15 and EBOV B1 with WFA, SBA or GNA used at a concentration of 10µg.mL^-1^ . Statistical significance was determined by one-way ANOVA followed by Sidak’s multiple comparison, p>0.05 (ns).

**Supplementary Figure 5.**
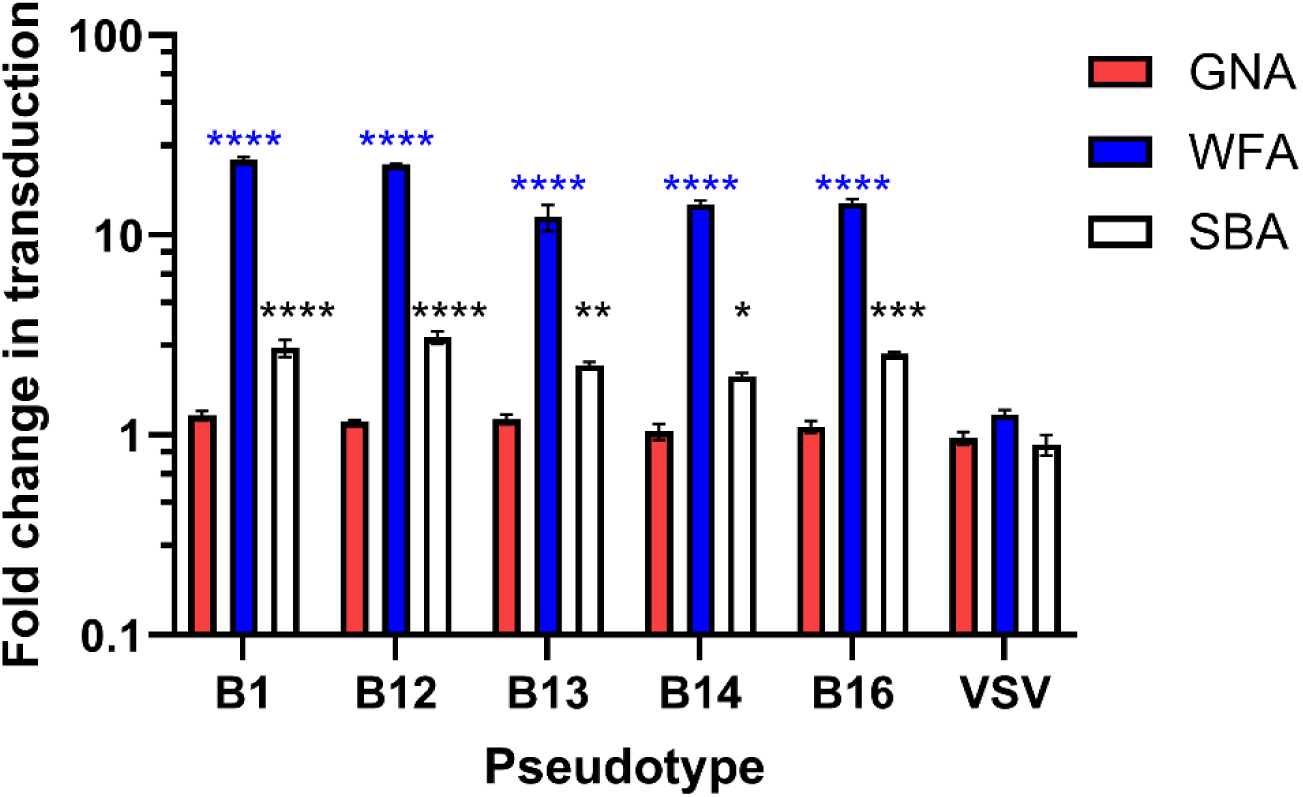
Enhancement of Ebola virus variants EBOV B1, B12, B13, B14, B16 with WFA, SBA or GNA used at a concentration of 10 µg.mL^-1^ . Pseudotypes possessing the VSV glycoprotein were used as a control in this experiment. Statistical significance was determined by one-way ANOVA followed by Dunnett’s multiple comparison test, p<0.05 (*), p<0.01 (**), p>0.0001 (****). Comparisons with p>0.05 are not labelled.

**Supplementary figure 6.**
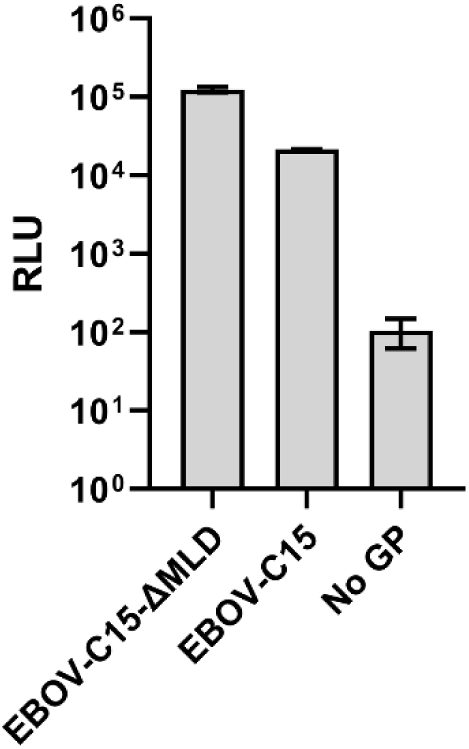
Transduction of HuH7 cells by pseudotypes bearing the EBOV-C15 and EBOV-C15-ΔMLD glycoproteins. Transduction was measured in relative light units (RLU) as a measure of luciferase reporter expression.

